# miRAssist: a context-aware, evidence integration framework for interpretable miRNA-target prioritization

**DOI:** 10.64898/2026.09.10.750642

**Authors:** Andrew Ring, Yaguang Xi

## Abstract

**Motivation:** MicroRNA-target interaction prediction remains challenging because many existing tools provide prediction scores or ranked candidate lists without making the supporting evidence easy to interpret or relate to a specific biological context.

**Results:** Here, we developed miRAssist, a context-aware evidence-integration framework for interpretable miRNA-target prioritization. miRAssist integrates six evidence families, including sequence complementarity, thermodynamic stability, sequence conservation, target-site accessibility, functional binding, and functional repression. A sequence-defined candidate universe was generated, resulting in 280,917 candidate interactions. Using miRTarBase-supported interactions as known-positive labels, six supervised scoring approaches were evaluated using a grouped train/test split by miRNA. Random forest showed the strongest performance and was selected. miRAssist also produced stronger known-positive enrichment than established miRNA-target prediction models in the evaluated benchmark. An LLM-assisted interface further supports natural-language database querying and evidence-grounded summarization of prioritized candidates.

**Availability and Implementation:** miRAssist is available as a web application at https://andy-ring-mirassist.share.connect.posit.cloud. The code is available at https://github.com/Andy-Ring/miRAssist

## 1 Introduction

First discovered in 1993, microRNAs (miRNAs) are small noncoding RNAs typically 20-25 nucleotides in length^1^. miRNAs regulate the expression of messenger RNA (mRNA) by binding to the 3’ untranslated region (3’ UTR)^2^. By doing so, miRNAs canonically repress the expression of mRNAs by blocking their translation^3^ or promoting their degradation^4^. In this role, miRNAs have been implicated in several biological processes^5^. Their dysregulation has also been shown to play a role in the pathogenesis of several diseases, including cancer^6^, cardiovascular disease^7^ and neurological disease^8^.

The role of miRNAs in biological processes has made them a focal point of research. An important aspect of that research has focused on understanding and predicting the interactions between miRNAs and mRNAs^9^. Through this work, some key features of the miRNA-mRNA interactions (MTIs) have been identified as having some predictive qualities^9,10^. The first 2-8 nucleotides in the miRNA sequence are known as the seed sequence, and a perfect Watson-Crick match of at least 6 nucleotides in that sequence is evidence of an MTI^5,11^. Seed matching, while not absolutely predictive, has been shown to be the foundation for MTIs and has been incorporated in most prediction tools^10,12^. Sequence conservation has also been shown to predict MTIs, as there is higher conservation of miRNA seed sequences compared to non-seed sequences^11,13,14^. This concept was used to develop the commonly used MTI prediction tool, TargetScan^15,16^. The next is free energy, the measurement of the energetic stability of the interaction between miRNA and mRNA^17^. This has also been incorporated into predictive models like RNAhybrid^17^ and miTarget^18^. However, strong seed-based evidence of interaction does not guarantee accessibility to that seed region^19^. mRNA secondary structure can make a seed region inaccessible and prevent miRNA binding^20^. Tools such as miRanda-mirSVR^21^ and DIANA-microT-CDS^22^ take site accessibility into account.

Many tools have sought to combine lines of evidence for MTIs using machine-learning models^21–24^. A common challenge faced is that many MTI prediction tools return a score or ranked list of targets without making the supporting evidence easy to interpret. This can make it difficult for users to understand why a target was prioritized or whether the evidence is relevant to their specific biological question. This is especially important because miRNA function is highly context-dependent^25^. A useful MTI prioritization tool ranks candidate interactions, explains the evidence supporting each candidate, and allows users to interpret those candidates in a biological context.

Here, we present miRAssist, a context-aware, evidence-integration framework for interpretable miRNA-target prioritization. miRAssist combines six evidence families: sequence complementarity, thermodynamic stability, sequence conservation, target-site accessibility, functional binding, and functional repression. These features are integrated into a transcript-level candidate database and used to train a backend random forest scoring model. The model was evaluated using held-out experimentally supported MTIs from miRTarBase and compared with established external prediction tools. To improve interpretability, miRAssist also includes a large language model (LLM)-based planner and synthesis layer. This layer allows users to ask natural-language questions and receive ranked candidates with evidence-grounded explanations.

## 2 Materials and Methods

### 2.1 Public Data Collection

Human mature miRNA sequences were downloaded from miRBase^26^. Human gene and transcript annotations were obtained from Gencode release 50 (Ensembl 116; GRCh38.p14)^27^. MANE Select transcripts were used as the primary transcript representation for candidate generation^28^. Experimentally supported miRNA-target interactions were obtained from miRTarBase^29^. TargetScanHuman8 predictions were used for sequence conservation^16^. miRNA-specific CLIP evidence was obtained from ENCORI^30^. Gene and miRNA expression data for breast invasive carcinoma (BRCA), colon adenocarcinoma (COAD), and prostate adenocarcinoma (PRAD) were obtained from the Cancer Genome Atlas (TCGA)^31^.

### 2.2 Candidate MTI Generation

The primary miRAssist candidate universe was generated independently of miRTarBase and external prediction databases. Mature human miRNAs were scanned against annotated 3′ UTR sequences from MANE Select transcripts. A miRNA-transcript pair was considered sequence eligible when the 3′ UTR contained at least one canonical 8mer, 7mer-m8, or 7mer-A1 site complementary to the mature miRNA. miRNA 5p and 3p strands were retained as distinct mature miRNAs, and versioned MANE transcript identifiers were preserved.

Because canonical seed matches alone produced a broad sequence-eligible universe, candidates were retained in the primary evidence table only when they also contained at least one selective source of independent support. A sequence-eligible pair was included when it had a directly mapped TargetScan prediction, a positive miRNA-specific CLIP-supported interaction, or expression evidence consistent with repression in at least one TCGA cancer type, defined as Spearman ρ ≤ −0.2 with Benjamini-Hochberg-adjusted q ≤ 0.05. RNAhybrid and RNAplfold evidence were not used to determine candidate eligibility. miRTarBase was also excluded from candidate generation. This procedure produced a primary candidate universe of 280,917 miRNA-transcript interactions.

### 2.3 Evidence Feature Construction

Evidence features were organized into six families: sequence complementarity, thermodynamic stability, sequence conservation, target-site accessibility, functional binding, and functional repression.

Sequence complementarity features were derived from canonical 8mer, 7mer- m8, and 7mer-A1 sites identified in MANE Select 3′ UTR sequences. Multiple qualifying sites within the same miRNA-transcript pair were summarized at the transcript level, including the best canonical site class and total number of qualifying sites.

Thermodynamic stability was calculated using RNAhybrid. RNAhybrid minimum free-energy values were calculated for candidate sites and summarized for each miRNA-transcript pair. These values were used as evidence features.

Target-site accessibility was estimated using RNAplfold from the ViennaRNA package. The best seed-region unpaired probability across qualifying sites was retained for each candidate. Failed or undefined calculations were retained as missing values rather than being assigned zero.

TargetScan predictions were mapped to sequence-defined candidates using transcript- and site-compatible mappings where available for sequence conservation evidence. TargetScan context++ scores and the externally supplied TargetScan context- score percentile were retained. Candidates without a valid TargetScan mapping remained in the evidence table when they qualified through another eligibility source.

Functional binding evidence was integrated at the miRNA-gene level using miRNA-specific ENCORI records. The number of supporting experiments and available CLIP score summaries were retained.

Functional repression evidence was generated separately for TCGA-BRCA, TCGA-COAD, and TCGA-PRAD. Spearman correlations were calculated between miRNA and mRNA expression for matched cancer-specific samples, and multiple-testing correction was performed separately within each cancer type using the Benjamini-Hochberg procedure. Negative correlations were interpreted as expression patterns consistent with miRNA-mediated repression.

### 2.4 Evidence Normalization and Summary Features

Raw evidence features were converted to percentile-based support measures using the same deterministic evidence-processing framework across the miRAssist evidence table. Internally derived percentiles were calculated globally across nonmissing candidate values using average ranks, with score direction standardized so that higher percentiles indicated stronger support. The externally supplied TargetScan context-score percentile was preserved rather than recalculated.

Evidence-family support percentiles were calculated from the available component percentiles within each family. Availability indicators distinguished the presence of valid evidence from favorable evidence. miRTarBase annotations and learned model scores were not included in any evidence-derived percentile or summary calculation.

### 2.5 Training and Testing Split

Candidates were grouped by mature miRNA using MIMAT accession and divided into training and testing sets using an 80/20 grouped split with a fixed random seed. All candidate targets for a given mature miRNA were assigned to only one partition. The training set contained 224,645 candidate interactions, including 2,067 known-positive interactions. The held-out test set contained 56,272 candidates, including 516 known-positive interactions. The test set contained 438 mature miRNAs, of which 65 had at least one aligned known-positive target.

### 2.6 Model Training and Evaluation

Six supervised scoring approaches were compared using the same training and testing partitions: logistic regression, XGBoost, random forest, support vector machine, multilayer perceptron, and naïve Bayes.^32,33^ Models were trained using known-positive miRTarBase interactions as the positive class and unlabeled candidate interactions as the background class. Therefore, the evaluation was interpreted as positive-unlabeled known-positive recovery rather than classification of experimentally confirmed positive and negative interactions.

Model discrimination was evaluated using the area under the receiver operating characteristic curve (AUROC) and precision-recall area under the curve (PR-AUC). The expected random PR-AUC was defined by the prevalence of known-positive interactions in the held-out evaluation set. Because miRAssist is intended as a prioritization system, ranking performance was additionally evaluated using Recall@3, Recall@5, Recall@10, and Recall@25 within each positive-bearing held-out miRNA group. The fraction of positive-bearing miRNAs with at least one known-positive interaction recovered within the top-ranked candidates was calculated separately.

### 2.7 Leave-One-Evidence-Family-Out Analysis

To estimate the incremental contribution of each evidence family, the random- forest model was retrained after removing one complete evidence family at a time.

Separate models were generated after omission of sequence complementarity, thermodynamic stability, sequence conservation, target-site accessibility, functional binding, and functional repression.

Each reduced model was evaluated on the same held-out test set using AUROC, PR-AUC, Recall@3, Recall@5, Recall@10, and Recall@25. Differences from the full random-forest model were evaluated using paired grouped-bootstrap resampling. These analyses were interpreted as measures of incremental predictive contribution within the current candidate universe rather than as measures of the biological importance of each evidence family.

### 2.8 External Model Comparison

The final miRAssist random-forest score was compared with TargetScan^15,16^, miRDB^23^, DIANA-microT^22^, miRanda^21^, and RNA22^24^ predictions.

TargetScanHuman 8 predictions were mapped using the TargetScan site mapping. miRDB v6.0 predictions were mapped from RefSeq transcript identifiers through NCBI human annotation release 110 to target genes and then to the corresponding MANE Select candidate. DIANA-microT 2023 predictions were mapped at the Ensembl gene level to the single corresponding MANE Select candidate. RNA22 v2 predictions were retained only when they mapped to the selected MANE transcript and a qualifying 3′ UTR site. miRanda v3.3a predictions were recomputed using the mature-miRNA sequences and MANE Select 3′ UTR sequences because a reproducible precomputed prediction set was not available.

External scores were transformed where necessary so that larger values consistently represented stronger predicted interaction support. For the full-universe comparison, candidates without a prediction from an external tool were assigned a sentinel score strictly below the weakest valid prediction from that tool, allowing coverage and discrimination to be evaluated jointly. Because prediction coverage differed substantially among external tools, pairwise common-coverage analyses were also performed in which miRAssist and each external method were evaluated only among candidates scored by that external method.

AUROC, PR-AUC, and Recall@3, Recall@5, Recall@10, and Recall@25 were calculated for each model. Confidence intervals and paired differences between miRAssist and each comparator were calculated using 1,000 grouped-bootstrap replicates by held-out miRNA.

### 2.9 LLM-Based Query Planning and Evidence Synthesis

The miRAssist application includes a natural-language layer for translating user questions into structured database retrievals and for summarizing the evidence associated with prioritized candidates. The LLM layer does not generate new evidence and does not independently validate biological interactions.

Query planning used a hybrid framework combining deterministic entity normalization with LLM-assisted semantic interpretation. Deterministic processing was used for miRNA and gene normalization, supported cancer contexts, evidence-family filters, novelty requests, result limits, and explicit query direction. Biological process and pathway intent were interpreted semantically and then resolved deterministically to supported pathway resources. Planner validation used 40 held-out natural-language queries, each evaluated across five repeated runs. The evaluation metrics included repeated natural language query benchmarks evaluating structured query agreement, retrieval critical agreement, retrieval equivalence, field-level accuracy, schema validity, execution success, ambiguity handling, and reproducibility.

Evidence synthesis was evaluated separately, where candidate identifiers, scores, ranks, and evidence statements were isolated by candidate before being supplied to the synthesis model. The synthesis layer was constrained to discuss only retrieved candidates and evidence contained in the supplied bundle. Evidence wording distinguished computational prediction, CLIP-supported binding, expression evidence consistent with repression, and known-positive annotation where available.

Synthesis validation used 40 held-out retrieval cases and three replicates. The evaluation metrics were candidate fidelity, numerical fidelity, rank fidelity, candidate-level evidence attribution, unsupported biological claims, fabrication, cross-candidate evidence transfer, and reproducibility.

### 2.10 Software

Data processing, database construction, model training, evaluation, and visualization were performed using Python. Machine learning models were trained and evaluated using scikit-learn^32^ and XGBoost^33^ packages. Additionally, RNAhybrid^34^ and RNAplfold^35^ were used to calculate the free energy and target-site accessibility, respectively. Evaluation metrics were calculated with scikit-learn. The front-end app implementation for miRAssist was developed using Python and Streamlit. The LLM models used for validation as well as the planner and synthesis steps of the miRAssist app, are OpenAI GPT 5.4-nano (planner) and GPT 5.4-mini (synthesis).

## 3 Results

### 3.1 Construction of miRAssist Database

A transcript-level miRNA-target candidate database was constructed by integrating evidence from key features of miRNA-target interaction. Each row represented one candidate miRNA-target interaction and was assigned a unique identifier to enable matching among the evidence table, held-out miRTarBase labels, and external model predictions.

Evidence features were grouped into 6 families (Table 1). Sequence complementarity features were constructed using miRNA and mRNA 3’UTR sequences to identify 8mer, 7mer-m8, and 7mer-A1 seed matches. Sequence conservation features were constructed from TargetScan data. Thermodynamic stability was calculated using RNAhybrid. Target-site accessibility features were generated using RNAplfold-based estimates of local unpaired probability. Functional binding features were generated using CLIP-seq hits collected from the ENCORI database. Functional repression features were generated using TCGA gene expression and miRNA expression data from 3 studies. In order to be included in the database, candidate MTIs must have had at least one 8mer, 7mer-m8, or 7mer-A1 site and evidence from sequence conservation, functional binding or functional repression. The resulting evidence table contained 280,917 potential MTIs (Figure 1A). A full list of features is described in Supplementary Table 1.

**Figure 1.**
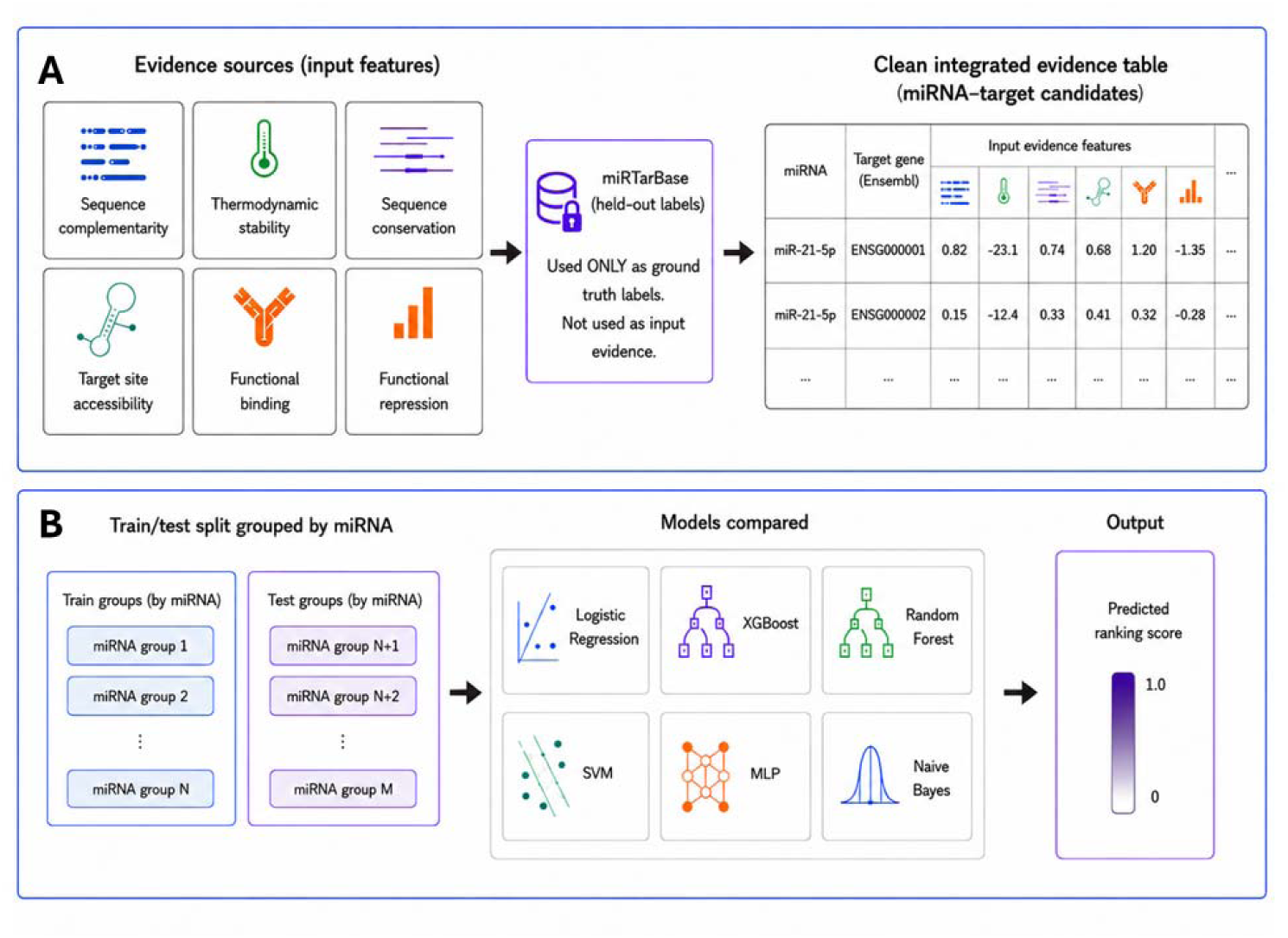
Overview of miRAssist evidence table construction and backend ranking model evaluation. **A.** miRAssist evidence table was constructed by combining features from 6 different MTI evidence families. Experimentally confirmed positive MTIs from miRTarBase were aligned but hidden for evaluation. **B.** To train and test a backend ML model to rank potential MTIs, each row of the evidence table was grouped by unique miRNA. miRNA groups were split 80/20 for training/testing. That data was used to train and evaluate 6 different ML model types. Each model was designed to output a score based on recovery of miRTarBase-supported known-positive interactions, which could be used to prioritize potential MTIs. Alt text: Two-panel workflow diagram of miRAssist database construction and model evaluation. Panel A shows six evidence families integrated into a miRNA–target evidence table: sequence complementarity, thermodynamic stability, sequence conservation, target-site accessibility, functional binding, and functional repression. miRTarBase interactions are reserved as held-out labels. Panel B shows miRNA-grouped training and testing sets used to compare six machine-learning approaches and produce candidate-ranking scores.

**Table 1.**
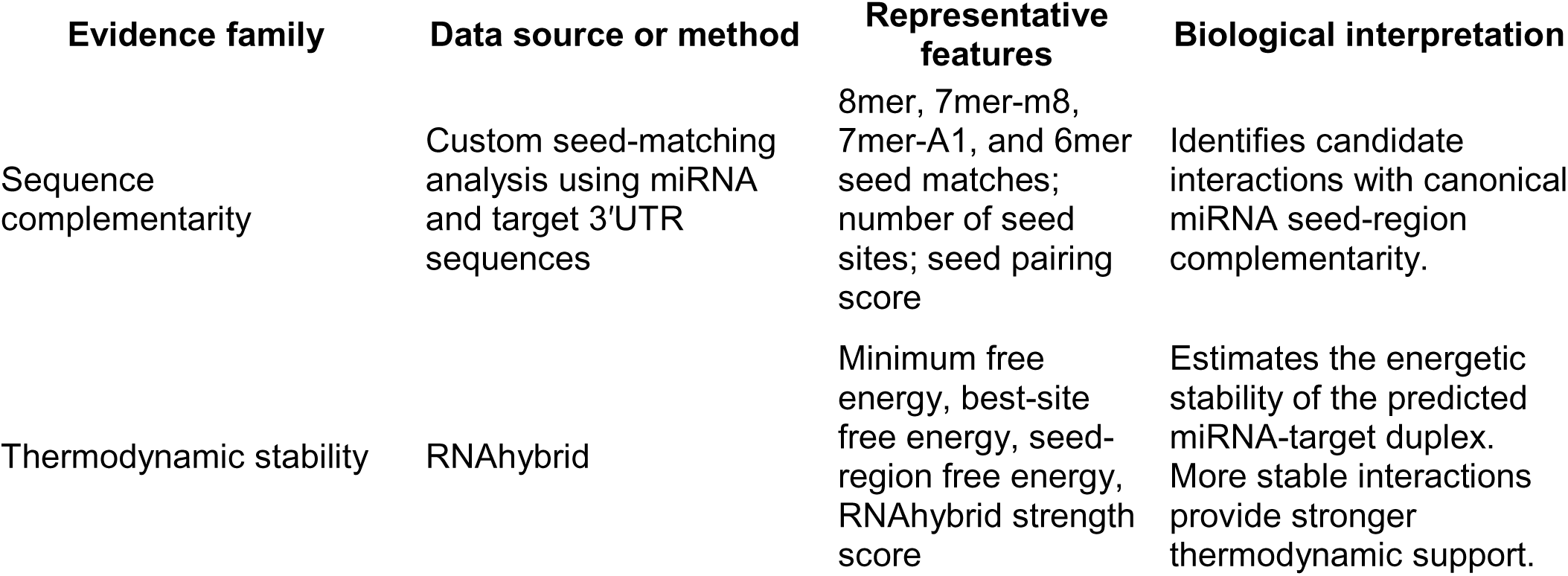
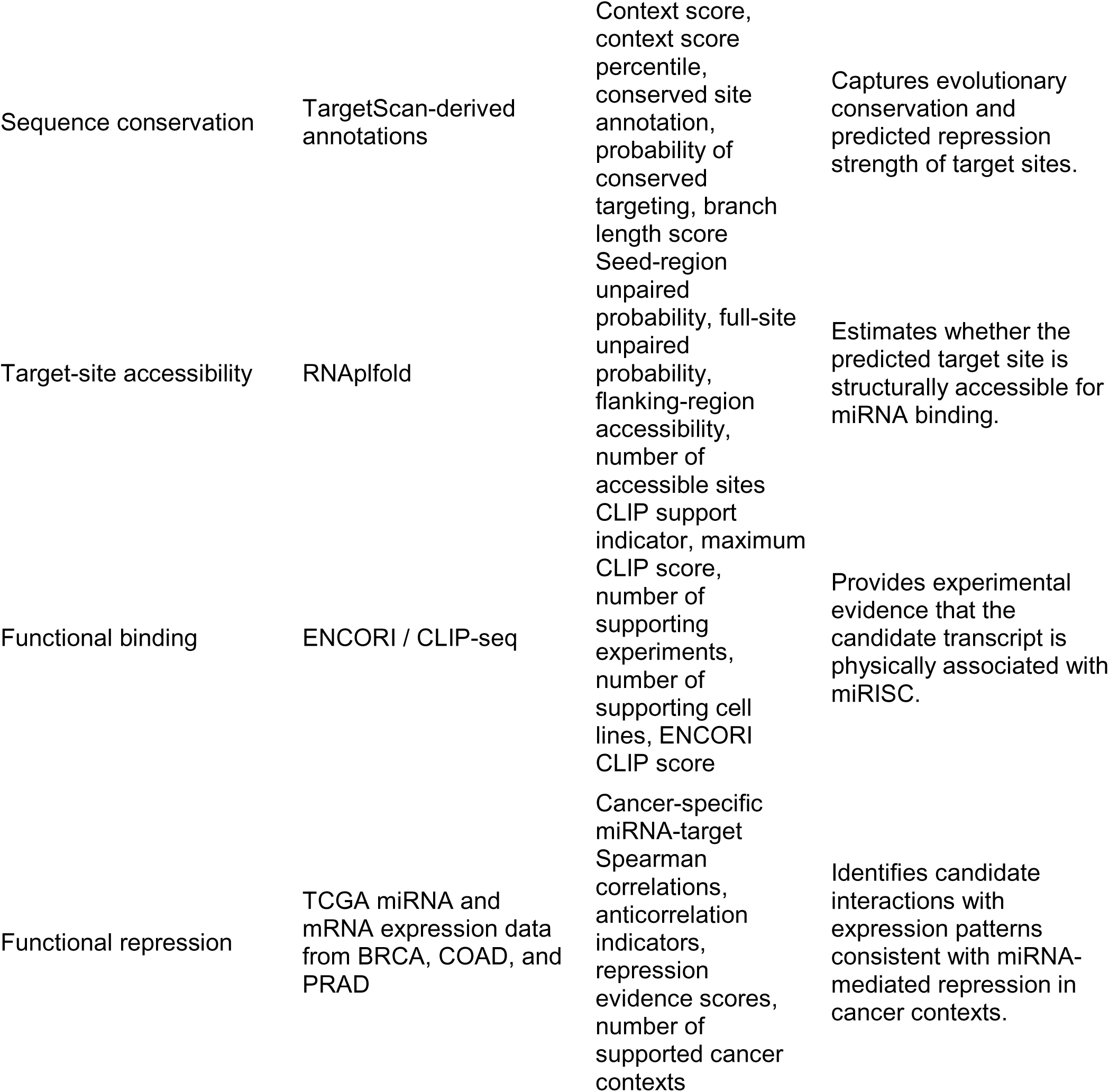
Overview of the evidence families integrated into the miRAssist database.

| Evidence family | Data source or method | Representative features | Biological interpretation |
| --- | --- | --- | --- |
| Sequence complementarity | Custom seed-matching analysis using miRNA and target 3'UTR sequences | 8mer, 7mer-m8, 7mer-A1, and 6mer seed matches; number of seed sites; seed pairing score | Identifies candidate interactions with canonical miRNA seed-region complementarity. |
| Thermodynamic stability | RNAhybrid | Minimum free energy, best-site free energy, seed-region free energy, RNAhybrid strength score | Estimates the energetic stability of the predicted miRNA-target duplex. More stable interactions provide stronger thermodynamic support. |
| Sequence conservation | TargetScan-derived annotations | Context score, context score percentile, conserved site annotation, probability of conserved targeting, branch length score | Captures evolutionary conservation and predicted repression strength of target sites. |
| Target-site accessibility | RNAplfold | Seed-region unpaired probability, full-site unpaired probability, flanking-region accessibility, number of accessible sites | Estimates whether the predicted target site is structurally accessible for miRNA binding. |
| Functional binding | ENCORI / CLIP-seq | CLIP support indicator, maximum CLIP score, number of supporting experiments, number of supporting cell lines, ENCORI CLIP score | Provides experimental evidence that the candidate transcript is physically associated with miRISC. |
| Functional repression | TCGA miRNA and mRNA expression data from BRCA, COAD, and PRAD | Cancer-specific miRNA-target Spearman correlations, anticorrelation indicators, repression evidence scores, number of supported cancer contexts | Identifies candidate interactions with expression patterns consistent with miRNA-mediated repression in cancer contexts. |

Additionally, experimentally supported miRNA-target interactions from miRTarBase were used during the evaluation. Strong evidence of interaction labels was joined to the evidence database by row-level candidate identifiers. miRTarBase labels were excluded from all model input features. The labels served as known-positive labels during evaluation. After alignment, there were 2,583 MTIs with strong evidence of interaction.

### 3.2 Creating training and testing sets with miRTarBase experimentally confirmed positives

To ensure the model provides top-ranked potential MTIs, we designed a backend machine learning model that prioritizes MTIs with evidence similar to that of confirmed positives in miRTarBase. To train and test the backend scoring model, we used a grouped train/test split by miRNA. Each unique miRNA was grouped with its potential targets. Then, the groups were split 80/20 for training and testing. This resulted in 1,751 miRNA groups in the training set, a total of 224,645 potential MTIs. 2,067 of those MTIs had a hidden label of strong evidence of interaction from miRTarBase. The testing set contained 438 miRNA groups, totaling 56,272 potential MTIs. 516 of those MTIs had the hidden label of strong evidence of interaction. Using this training system, the backend model should prioritize MTIs with features similar to experimentally confirmed positives (Figure 1B).

### 3.3 Random forest outperforms other backend scoring models

We compared several backend ranking approaches using the same blinded evidence table and the same training/testing groups. The models tested included XGBoost, logistic regression, random forest, support vector machine (SVM), multilayer perceptron (MLP), and naïve Bayes. The models’ performance was assessed using AUROC, PR-AUC, and Recall@K on the testing set.

Random forest showed the strongest performance of all models tested. On the held-out testing set, it achieved an AUROC of 0.846 and a PR-AUC of 0.135. Because the test set contained 516 miRTarBase-supported interactions among 56,272 candidate miRNA–target interactions (MTIs), the positive-label prevalence was approximately 0.92%, corresponding to an expected random PR-AUC baseline of approximately 0.0092. Thus, the miRAssist random forest score achieved a PR-AUC approximately 14-fold higher than the random baseline, indicating substantial enrichment of known-positive interactions over random ranking. XGBoost showed a similar AUROC of 0.840 but a reduced PR-AUC of 0.104. Logistic regression, SVM, and MLP showed slightly lower performance, with AUROCs of 0.817, 0.805, and 0.821 and PR-AUCs of 0.085, 0.080, and 0.106, respectively. Naïve Bayes finished last with an AUROC of 0.752 and a PR-AUC of 0.028 (Figure 2A, B).

**Figure 2.**
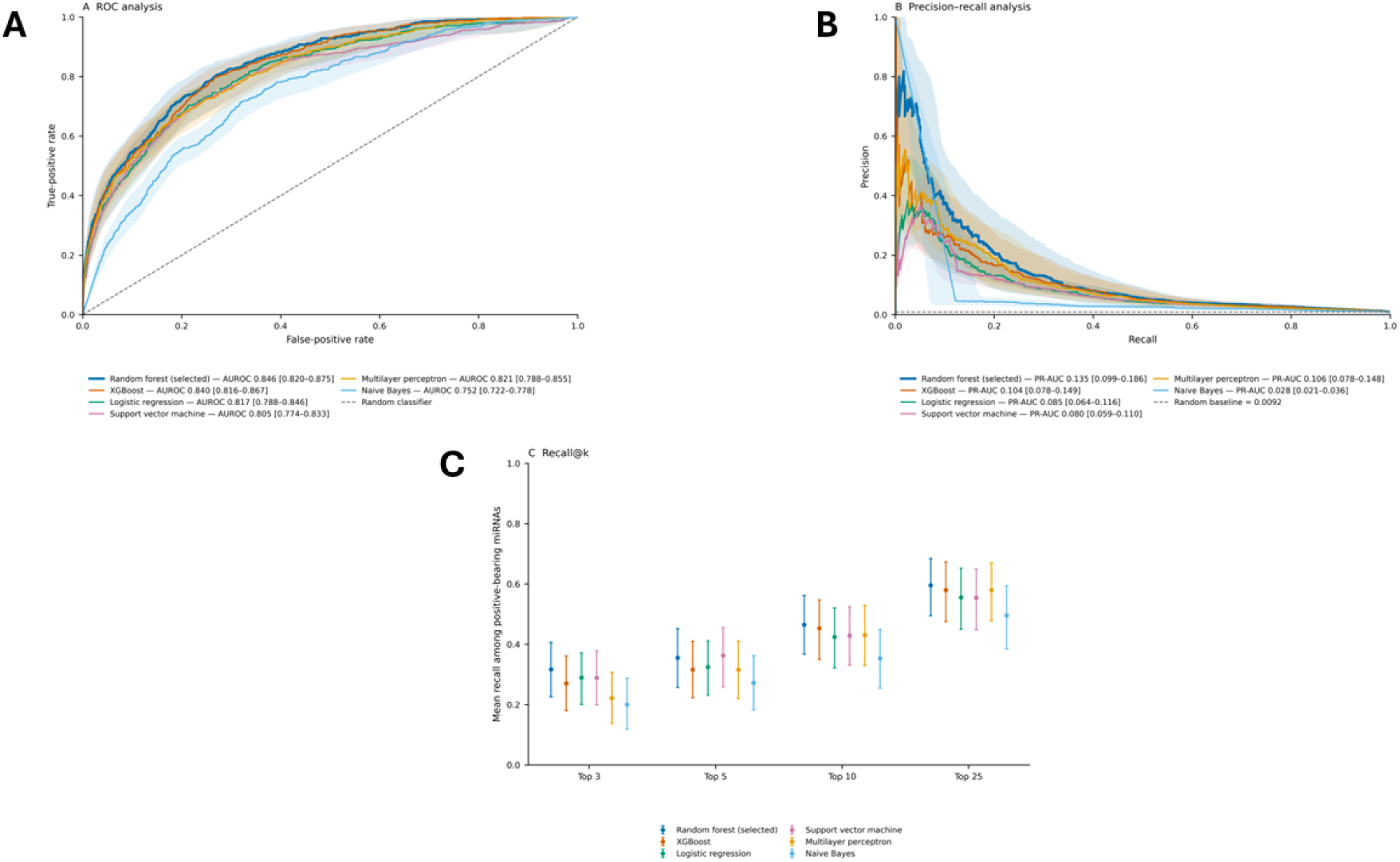
Comparison of different machine learning models for ranking potential MTIs. **A.** Receiver operating characteristic curve, **B.** Precision-recall curve, and **C.** Recall@K analysis of potential machine learning models’ performance on the evaluation set. Alt text: Three-panel comparison of six machine-learning models on the held-out candidate set. Panels A and B compare receiver operating characteristic and precision– recall curves for random forest, XGBoost, logistic regression, support vector machine, multilayer perceptron, and naïve Bayes. Random forest achieves the highest AUROC and PR-AUC. Panel C shows mean Recall at the top 3, 5, 10, and 25 candidates, where random forest gives the strongest top-ranked recovery.

To evaluate backend performance in a manner that more directly reflects the intended use of miRAssist as a candidate-prioritization tool, we compared mean Recall@K across the 65 positive-bearing held-out miRNA groups. Random forest showed the strongest overall recovery, with mean Recall@3, Recall@5, Recall@10, and Recall@25 values of 0.317, 0.355, 0.465, and 0.596, respectively. XGBoost showed the next strongest performance at most cutoffs, with corresponding recall values of 0.270, 0.316, 0.453, and 0.580, while logistic regression achieved 0.289, 0.324, 0.424, and 0.555. Support vector machine performance was similar, with Recall@3/5/10/25 values of 0.289, 0.362, 0.428, and 0.554, and the multilayer perceptron achieved 0.222, 0.315, 0.430, and 0.580. Naïve Bayes showed the weakest top-ranked recovery, with recall values of 0.200, 0.272, 0.353, and 0.495. Random forest also recovered at least one known-positive interaction within the top 10 candidates for 84.6% of positive-bearing held-out miRNAs and within the top 25 for 90.8% (Figure 2C).

### 3.4 Functional evidence makes the largest contribution to model performance

To assess the contribution of each evidence family, we performed a leave-one-family-out analysis in which each evidence family was removed in turn and the random- forest model was retrained and evaluated on the same held-out test set. The full model achieved an AUROC of 0.846 and a PR-AUC of 0.135. Removing functional binding evidence produced the largest decline in PR-AUC, which decreased to 0.060, corresponding to a reduction of 0.075. Removing functional repression evidence also substantially reduced performance, decreasing AUROC from 0.846 to 0.752 and PR-AUC from 0.135 to 0.069, a PR-AUC reduction of 0.066. Sequence conservation evidence made a smaller but measurable contribution, with PR-AUC decreasing to 0.113 after its removal. Removing target-site accessibility reduced PR-AUC to 0.118, while removal of sequence complementarity or thermodynamic stability had comparatively little effect, both yielding PR-AUC values of 0.127 (Figure 3A, B).

**Figure 3.**
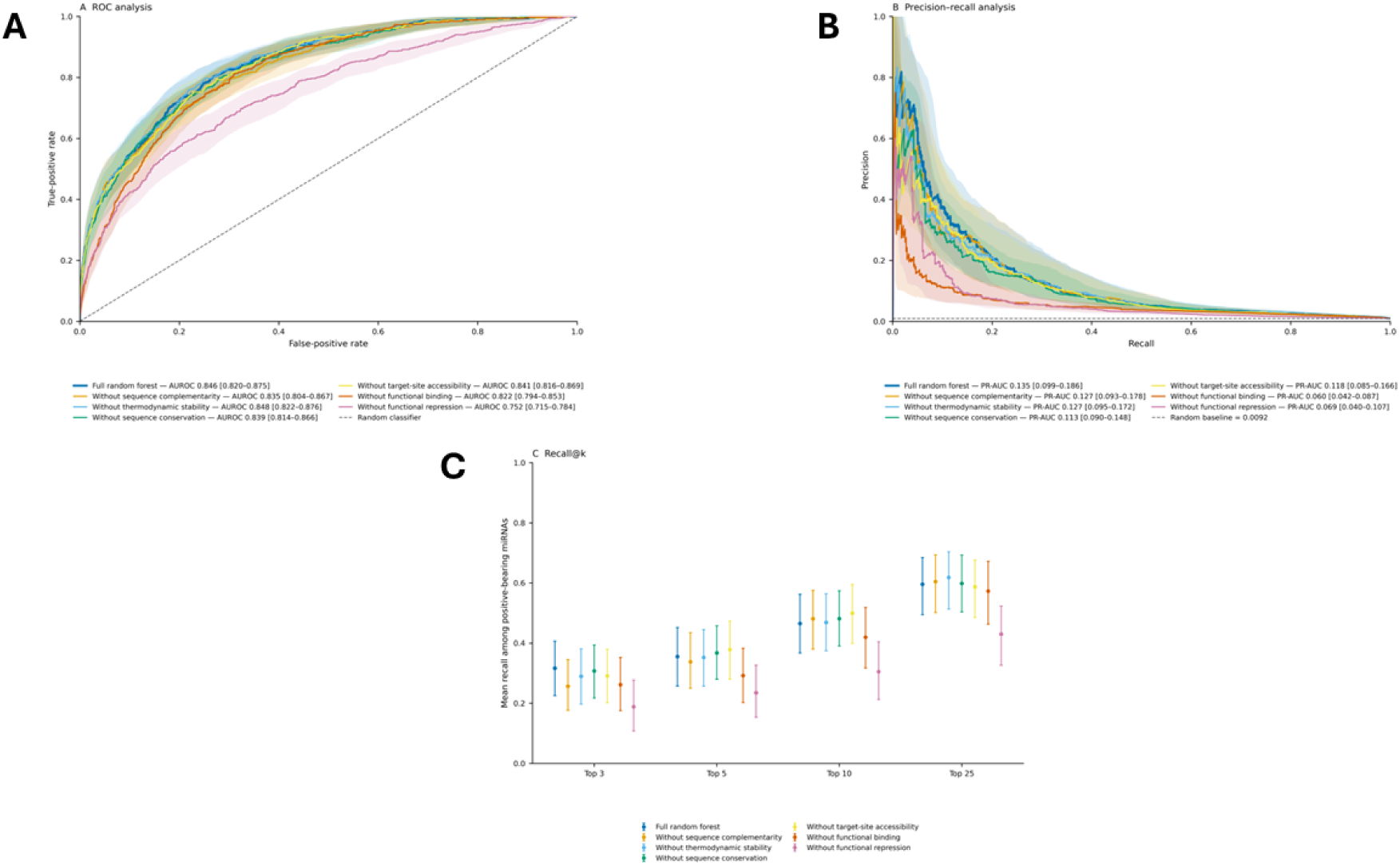
Evaluation of the contribution of each evidence family using leave-one-out analysis. **A.** Receiver operating characteristic curve, **B.** Precision-recall curve and **C.** Recall@K analysis of random forest models trained on remaining evidence. Alt text: Three-panel leave-one-evidence-family-out analysis of the random-forest model. Panels A and B compare receiver operating characteristic and precision–recall curves after removing each evidence family. Removing functional repression produces the largest reduction in AUROC, while removing functional binding produces the largest reduction in PR-AUC. Panel C shows that removing functional repression causes the largest decrease in Recall at the evaluated ranking cutoffs.

We next examined whether removal of individual evidence families affected recovery of miRTarBase-supported interactions near the top of each miRNA-specific candidate ranking. The full random-forest model achieved mean Recall@3, Recall@5, Recall@10, and Recall@25 values of 0.317, 0.355, 0.465, and 0.596, respectively.

Removal of functional repression evidence produced the largest and most consistent loss in top-ranked recovery, reducing Recall@3/5/10/25 to 0.188, 0.235, 0.305, and 0.430. Removing functional binding also reduced recovery, with corresponding recall values of 0.261, 0.292, 0.419, and 0.573. In contrast, omission of sequence complementarity, thermodynamic stability, sequence conservation, or target-site accessibility produced comparatively modest changes in Recall@K, and in some cases small increases at individual cutoffs. These ranking-based results were consistent with the PR-AUC analysis and further indicate that functional repression and functional binding evidence contribute most strongly to concentrating known-positive interactions near the top of the prioritized candidate lists (Figure 3C).

### 3.5 Integrated miRAssist scoring improves known-positive recovery relative to individual external prediction tools

We next compared the final miRAssist random forest model with five established miRNA-target prediction tools: TargetScan, miRDB, DIANA-microT, miRanda, and RNA22. Because the external tools differed substantially in candidate coverage, performance was evaluated both across the complete held-out test set and on pairwise common-coverage subsets. On the full held-out set, miRAssist achieved an AUROC of 0.846 and a PR-AUC of 0.135, compared with AUROC/PR-AUC values of 0.611/0.017 for TargetScan, 0.722/0.028 for miRDB, 0.711/0.032 for DIANA-microT, 0.584/0.012 for miRanda, and 0.480/0.009 for RNA22 (Figure 4A, B). External-tool coverage ranged from 39.5% of held-out candidates for TargetScan to 98.9% for DIANA-microT. To separate ranking performance from incomplete prediction coverage, we additionally compared miRAssist and each external tool only among candidates scored by that tool. miRAssist retained substantially higher PR-AUC on every common-coverage subset, achieving PR-AUC values of 0.202 versus 0.024 for TargetScan, 0.155 versus 0.034 for miRDB, 0.136 versus 0.032 for DIANA-microT, 0.138 versus 0.012 for miRanda, and 0.155 versus 0.009 for RNA22 (Figure S1).

**Figure 4.**
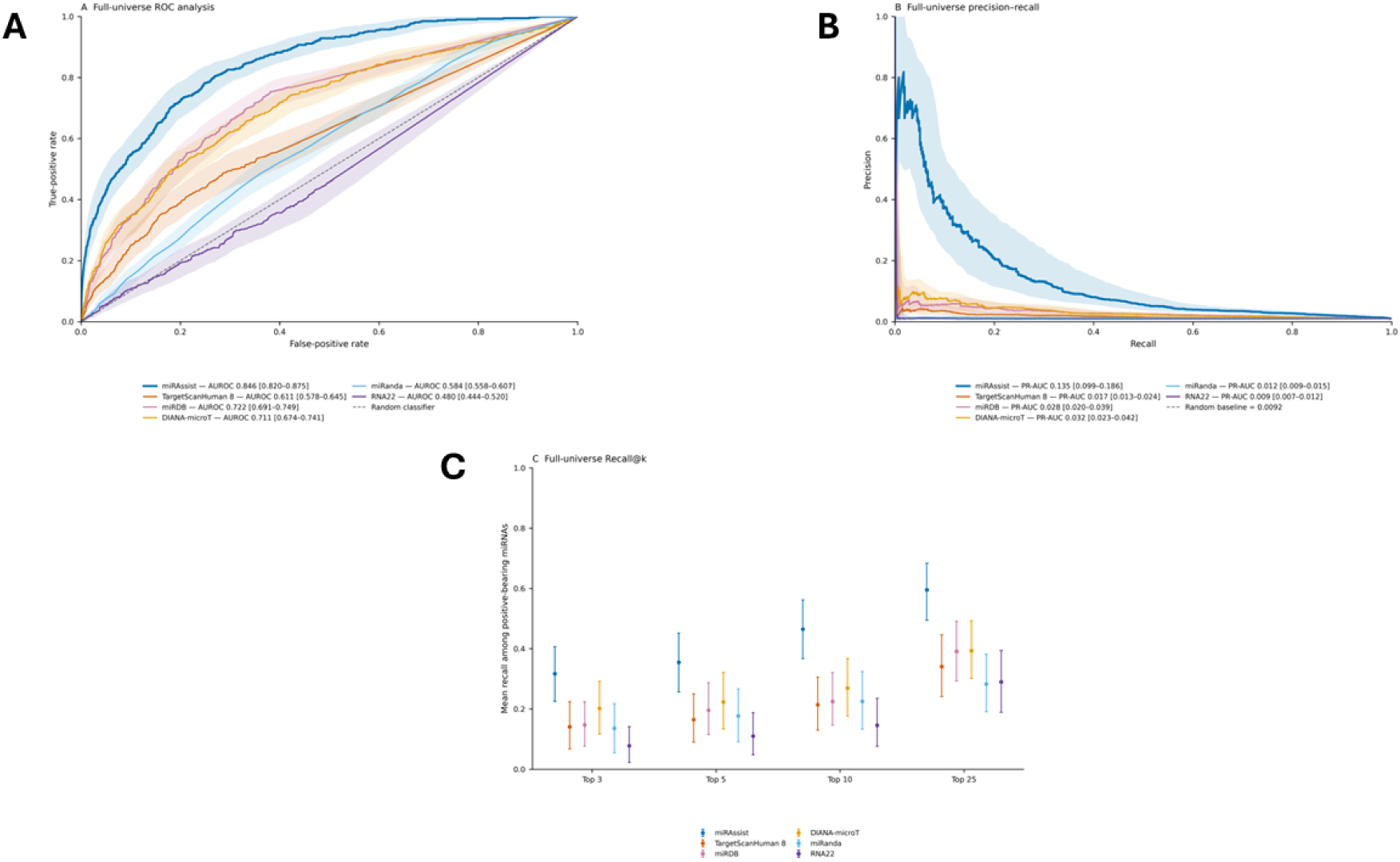
Comparison of external MTI prediction models with miRAssist. **A.** Receiver operating characteristic curve and **B.** Precision-recall curve, and **C.** Recall@K analysis of external MTI prediction models’ performance on the evaluation set. Alt text: Three-panel comparison of miRAssist with TargetScan, miRDB, DIANA-microT, miRanda, and RNA22 across the complete held-out candidate set. Panels A and B show that miRAssist has the highest AUROC and PR-AUC. Panel C shows that miRAssist also achieves the highest mean Recall at the top 3, 5, 10, and 25 candidates.

Ranking-based evaluation produced a similar pattern. The miRAssist random- forest model achieved mean Recall@3, Recall@5, Recall@10, and Recall@25 values of 0.317, 0.355, 0.465, and 0.596, respectively, across the 65 positive-bearing held-out miRNAs. Among the external tools, DIANA-microT showed the strongest early recovery, with Recall@3/5/10/25 values of 0.202, 0.223, 0.269, and 0.393. miRDB achieved corresponding values of 0.147, 0.196, 0.225, and 0.392, while TargetScan achieved 0.141, 0.165, 0.214, and 0.341. miRanda produced Recall@3/5/10/25 values of 0.136, 0.177, 0.225, and 0.282, and RNA22 showed the lowest top-ranked recovery, with values of 0.078, 0.110, 0.146, and 0.290 (Figure 4C).

### 3.6 miRAssist supports context-aware interpretation of natural language queries using large language models

To support natural-language interaction with miRAssist, we implemented an LLM-assisted planner and synthesis layer around the miRAssist evidence database and random-forest prioritization model. The planner combines deterministic entity and context normalization with LLM-based semantic interpretation to translate user questions into structured database queries. In a held-out benchmark of 40 natural-language queries, the planner produced schema-valid and executable queries in 100% of cases, with 100% reproducibility and no hallucinated biological entities. Although exact structured-query agreement was 57.5%, the expected candidate retrieval was reproduced exactly for 92.5% of queries, with a mean candidate-set Jaccard similarity of 93.5% and 85.0% agreement for the top-ranked candidate (Figure S2). This distinction reflects cases in which different structured representations produced equivalent or near-equivalent database retrievals.

The synthesis layer summarizes the retrieved candidates using their miRAssist scores, ranks, and supporting evidence while restricting interpretation to information contained in the retrieved evidence bundle (Figure 5A). In held-out synthesis validation, candidate identity, numerical values, and ranks were reproduced with 100% fidelity, and evidence attribution achieved 100% precision and 89.7% recall. All evaluated responses were classified as grounded under the prespecified criteria, with no unsupported biological claims, score-as-probability statements, or database scope violations observed (Figure S3). Together, these results support the use of the LLM layer for context-aware database retrieval and evidence-grounded summarization rather than as an independent source of biological evidence or validation.

**Figure 5.**
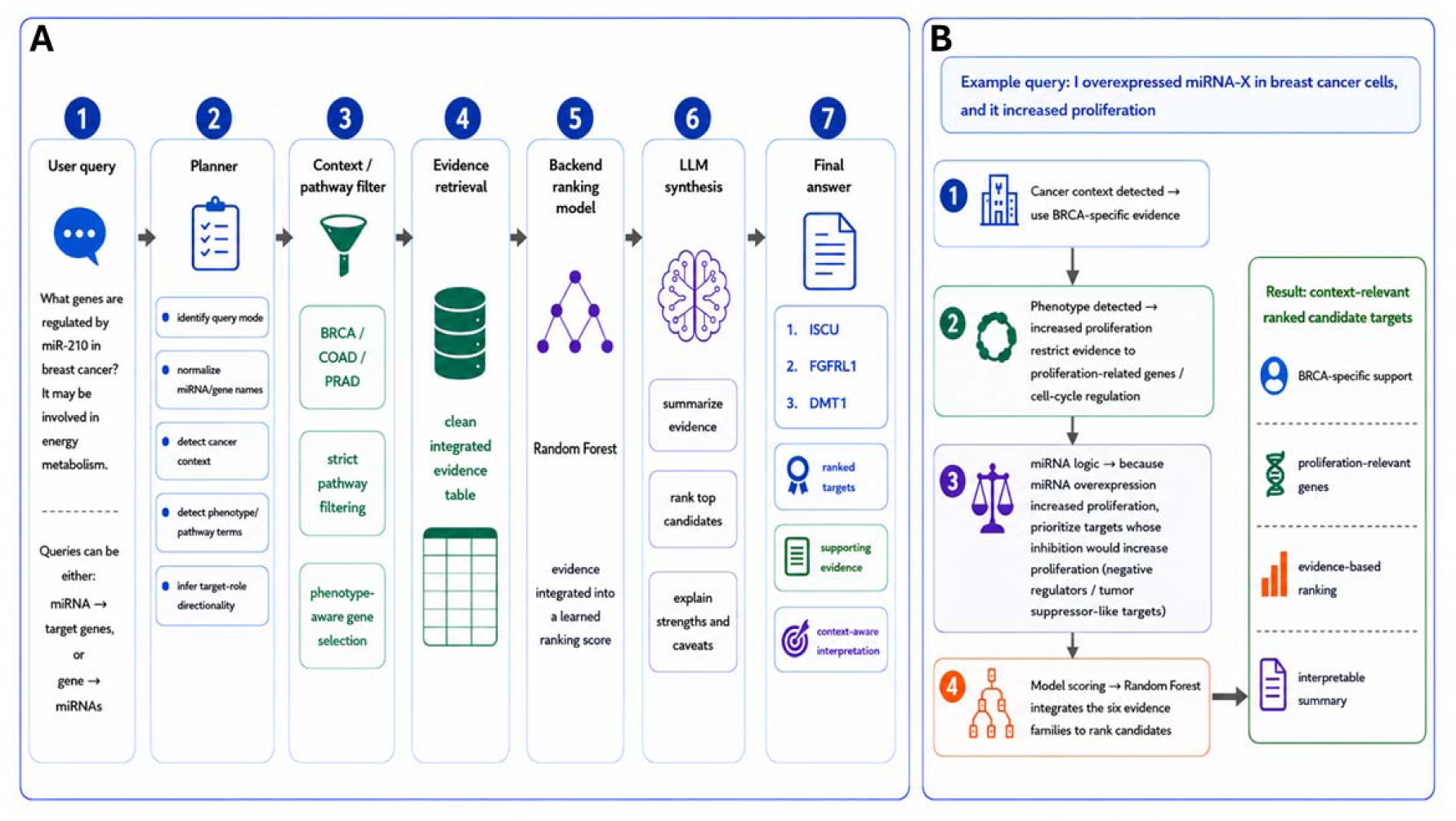
Overview of final miRAssist model. **A.** Flow of a user query through miRAssist. In brief, users enter a natural-language query into miRAssist, and the planner layer combines deterministic normalization with LLM-assisted semantic interpretation to construct a structured database query. Then it applies any relevant filtering constraints and executes the database retrieval. Then the database returns candidates ranked by their random forest model score, and the synthesis LLM writes a description of the top-ranked candidates and their evidence. **B.** Summary of how the planner LLM takes context into account when generating a database query. It can take cancer context, biological process, and implied directionality into account when synthesizing the database query. Alt text: Two-panel diagram of the miRAssist natural-language interface. Panel A follows a user question through query planning, cancer-context and pathway filtering, evidence retrieval, random-forest ranking, LLM-based synthesis, and generation of an evidence-grounded answer. Panel B illustrates how an example breast cancer query is interpreted using cancer context, proliferation-related phenotype information, and miRNA directionality. These elements are used to produce context-relevant ranked targets with supporting evidence.

## 4 Discussion

Substantial progress has been made in the prediction of miRNA target interactions^9,12^. However, despite these advances, predictive models still rely heavily on a single source of evidence or combine evidence in an uninterpretable way. To address this, we developed miRAssist, a context-aware framework for prioritizing miRNA-target interactions through evidence integration and interpretable synthesis. miRAssist combines six evidence families into a transcript-level candidate database and uses a trained random forest backend model to rank potential MTIs. The ranked results are then passed to an LLM-based layer that helps users ask natural-language questions and interpret the evidence supporting each candidate. We aim for miRAssist to empower researchers during the discovery and planning stages of miRNA research by streamlining MTI search and prioritization.

One notable result of the leave one evidence family out analysis was the disproportionate contribution of specific evidence families to the random forest model’s predictive power. We found that both functional binding and functional repression data contributed strongly to the model. This finding is biologically plausible because these evidence types capture experimentally observed binding or expression patterns associated with miRNA–target regulation as opposed to computational prediction. The strong contribution of these two evidence families also shows that there is an element dictating the interaction that is not captured by the sequence-based evidence families. It should be noted that though sequence complementarity, thermodynamic stability, and target site accessibility did not contribute meaningfully to this model, we cannot determine if that is due to technical or biological reasons.

During evaluation, we showed that miRAssist outperforms comparable external models. The task each model was asked to complete was to prioritize potential MTIs, with the goal of placing miRTarBase-confirmed positives near the top. miRAssist performed significantly better at this than any of the external models in the full held-out test set evaluation. Additionally, using pairwise common coverage test sets, miRAssist was compared to an external model using only the held-out MTIs that were present in the external model’s available data; miRAssist also outperformed each model. This evaluation was complicated by a few factors. First, there are very few miRTarBase positives among all potential MTIs. This causes precision metrics to appear lower than those reported in other publications. Additionally, since miRTarBase is a collection of experimentally confirmed positives, it is not an exhaustive list of true positive interactions, and there are no confirmed true negatives in the dataset. This problem has been documented^36^ and is persistent across all MTI prediction models. The enhanced interpretability of miRAssist aims to address this by combining enrichment of confirmed positives and other MTIs with similar features across the 6 evidence families, with clearly presented evidence for the researcher to use when determining which candidate to pursue.

## Authors’ contributions

A.R. and Y.X. were responsible for conceptualization of the research goals and aims.

A.R. developed the database and software, performed evaluation and benchmarking.

A.R. and Y.X. wrote the manuscript. Y.X. managed project administration and funding acquisition.

## Conflicts of interest

The authors declare no conflicts of interest.

## Funding

This work was supported in part by the National Institutes of Health (NIH) grants R01CA271533, R01CA260698, and R01CA275089, and by the U.S. Department of Veterans Affairs (VA), Biomedical Laboratory Research and Development Service, VA Merit Review Award I01 BX005094, all awarded to Y.X.

## Data and code availability

The code and data used for miRAssist, and the evaluation are available on GitHub at https://github.com/Andy-Ring/miRAssist. The application itself is free to use and is available at https://andy-ring-mirassist.share.connect.posit.cloud.

## Acknowledgements

The authors would like to thank the Georgia Advanced Computing Resource Center (GACRC) at the University of Georgia for providing computational resources.

## Notes

https://github.com/Andy-Ring/miRAssist

https://andy-ring-mirassist.share.connect.posit.cloud

### Competing Interest Statement

The authors have declared no competing interest.

