## Supplementary material for "miRAssist: a context-aware, evidence integration framework for interpretable miRNA-target prioritization": S_Fig

**Supplementary Figures**

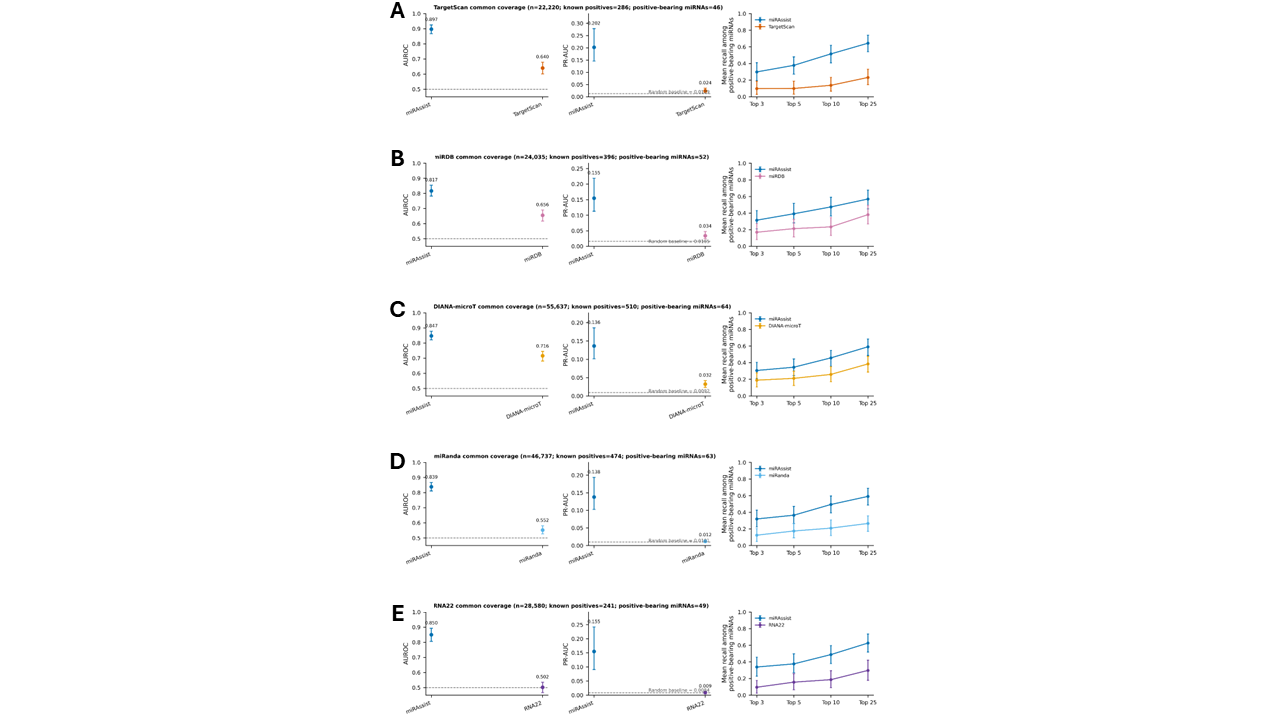

**Figure S1. Comparisons of miRAssist to external models using common coverage in the held-out miRNA set.** miRAssist was compared to **A.** TargetScan, **B.** miRDB, **C.** DIANA-microT, **D.** miRanda, **E.** RNA22.

Alt text: Five-row comparison of miRAssist with TargetScan, miRDB, DIANA-microT, miRanda, and RNA22 on separate pairwise common-coverage subsets. Each row presents AUROC, PR-AUC, and mean Recall at the top 3, 5, 10, and 25 candidates. miRAssist outperforms the corresponding external method across all five shared-coverage comparisons.

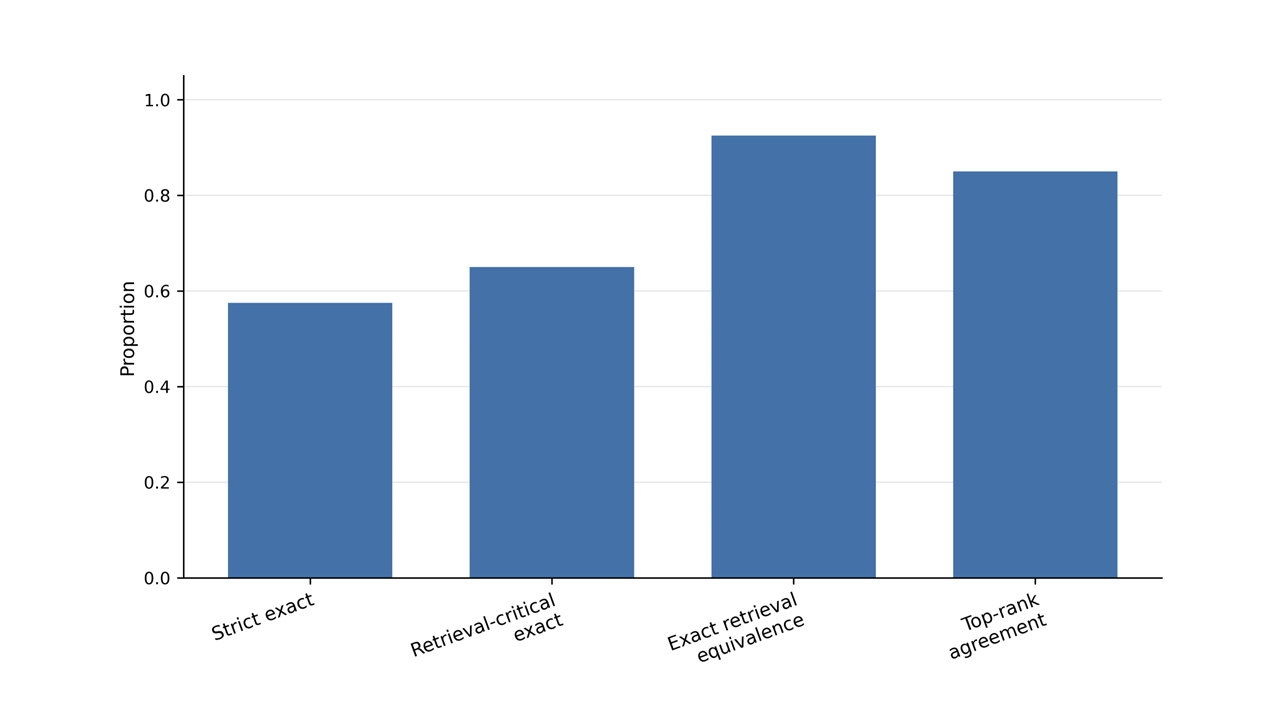

**Figure S2. Validation of the miRAssist natural-language planner on held-out queries.** Planner performance was evaluated using strict structured-query agreement, retrieval-critical agreement, exact retrieval equivalence, and agreement of the top-ranked candidate.

Alt text: Bar chart summarizing validation of the miRAssist natural-language query planner. Strict structured-query agreement is the lowest of the four measures, while exact retrieval equivalence exceeds 90%. Retrieval-critical agreement and agreement on the top-ranked candidate fall between these values. The results indicate that different structured queries frequently produce equivalent or closely matching retrieval results.

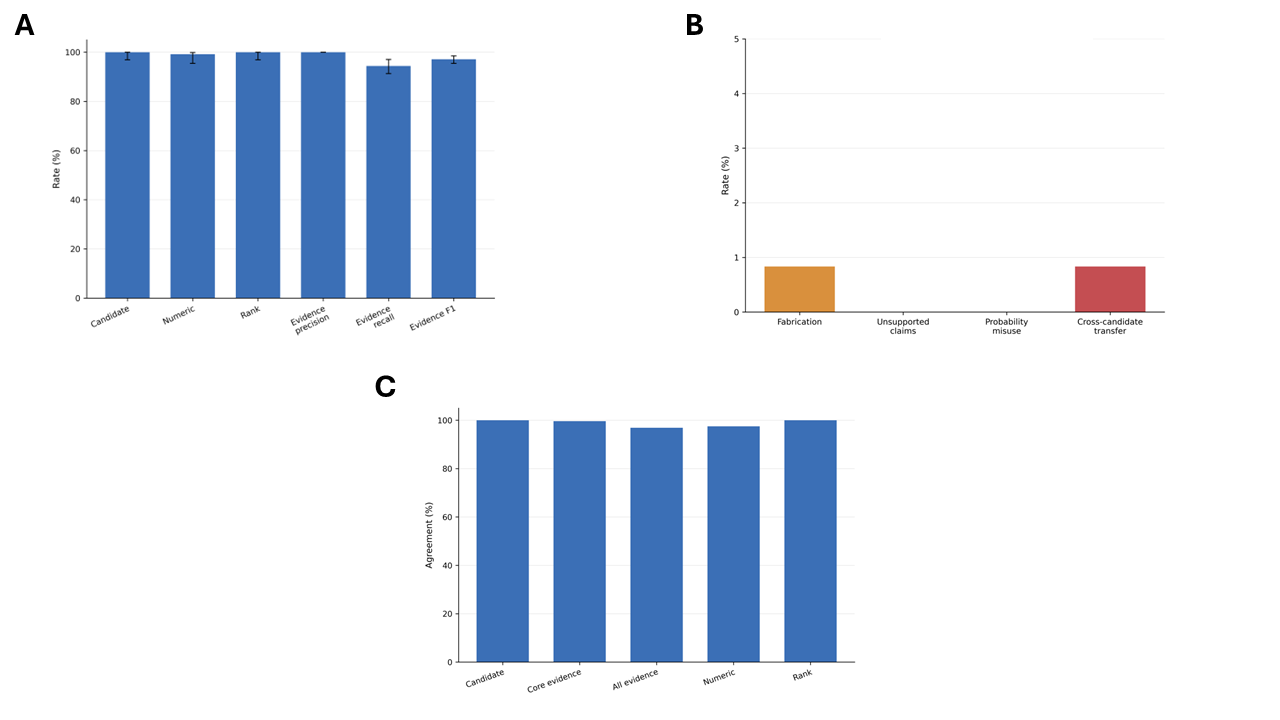

**Figure S3. Validation of the miRAssist synthesis layer on held-out retrieval bundles.** **A.** Grounding fidelity for candidate identity, numerical values, ranks, and evidence attribution, **B.** error rates for fabrication, unsupported claims, score-probability misuse, and cross-candidate evidence transfer, **C.** reproducibility across three synthesis replicates.

Alt text: Three-panel evaluation of the miRAssist evidence-synthesis layer. Panel A shows near-perfect fidelity for candidate identity, numerical values, ranks, and evidence attribution. Panel B shows very low error rates, with no unsupported-claim or probability-misuse errors and small nonzero rates of fabrication and cross-candidate evidence transfer. Panel C shows high reproducibility across replicates for candidate identity, evidence content, numerical values, and ranks.
